# SARS-CoV-2 mutations affect proteasome processing to alter CD8^+^ T cell responses

**DOI:** 10.1101/2022.04.08.487623

**Authors:** Dannielle Wellington, Zixi Yin, Zhanru Yu, Raphael Heilig, Simon Davis, Roman Fischer, Suet Ling Felce, Philip Hublitz, Ryan Beveridge, Danning Dong, Guihai Liu, Xuan Yao, Yanchun Peng, Benedikt M Kessler, Tao Dong

## Abstract

Viral CD8^+^ epitopes are generated by the cellular turnover of viral proteins, predominantly by the proteasome. Mutations located within viral epitopes can result in escape from memory T cells but the contribution of mutations in flanking regions of epitopes in SARS-CoV-2 has not been investigated. Focusing on two of the most dominant SARS-CoV-2 nucleoprotein CD8^+^ epitopes, we identified mutations in epitope flanking regions and investigated the contribution of these mutations to antigen processing and T cell activation using SARS-CoV-2 nucleoprotein transduced B cell lines and *in vitro* proteasomal processing of peptides. We found that decreased NP_9-17_-B*27:05 CD8^+^ T cell responses to the NP-Q7K mutation correlated with lower epitope surface expression, likely due to a lack of efficient epitope production by the proteasome, suggesting immune escape caused by this mutation. In contrast, NP-P6L and NP-D103N/Y mutations flanking the NP_9-17_-B*27:05 and NP_105-113_-B*07:02 epitopes, respectively, increased CD8^+^ T cell responses associated with enhanced epitope production by the proteasome. Our results provide evidence that SARS-CoV-2 mutations outside the epitope could have a significant impact on antigen processing and presentation, thereby contributing to escape from immunodominant T cell responses. Alternatively, mutations could enhance antigen processing and efficacy of T cell recognition, opening new avenues for improving future vaccine designs.

**One Sentence Summary:** Natural mutations in the flanking regions of known immunodominant SARS-CoV-2 nucleoprotein epitopes can decrease CD8^+^ T cell responses leading to partial escape.

## INTRODUCTION

Defining CD8^+^ T cell responses to SARS-CoV-2 has been a priority since the virus first emerged in 2019 as these responses are instrumental in clearing viral infections (*1*). Multiple immunodominant CD8^+^ T cell epitopes have been identified (*2-7*). Generation of CD8 epitopes initially requires antigen processing through the proteasome, a multi-subunit cytosolic machinery that can degrade intracellular proteins following ubiquitination. The proteasome is composed of a catalytic core particle (20s) and one or two terminal 19s regulatory particles. The 20s catalytic core is capable of cleaving at the C-terminal side of acidic, basic and hydrophobic amino acid residues (*8*). As part of this proteolytic process, a small fraction of peptide fragments will contain correctly cleaved C-termini of CD8 epitopes, whereas the N-terminus will generally require further trimming by aminopeptidases (*9*).

Further trimming of peptides can occur in the cytoplasm by various peptidases or, more commonly, in the endoplasmic reticulum (ER) following translocation of peptides from the cytosol via TAP. An essential peptidase responsible for trimming peptides in the ER is ERAP1; an aminopeptidase optimised to generate peptides of 8-10 amino acids for binding to MHC Class I molecules (*10*). ERAP1 activity varies depending on the N-terminal amino acid with little ability to cleave after proline residues and high ability to cleave after hydrophobic amino acids (*11*).

Once cleaved to optimal length, peptides will bind to MHC Class I molecules and those with sequences that bind strongly into the peptide-binding groove, so-called high affinity peptides, will induce transport of peptide-MHC complexes to the cell surface for activation of CD8^+^ T cells (*12*). Thus, the sequence of the peptides is important for ensuring strong binding to MHC, and mutations within these sequences can alter the affinity of this interaction. Using exogenous peptide assays, the effect of mutations within epitope sequences have been previously shown to lead to escape from viral T cell responses in various settings (*13-15*) but also in the context of SARS-CoV-2 (*16*).

In addition, components of the antigen processing pathway have preferences for amino acid sequences for their optimal processing as discussed above. Indeed, previous studies looking into the effect of viral mutations in the flanking regions of epitopes on antigen processing of HIV-1 found evidence of disruption of proteasomal cleavage (*17*) and endoplasmic reticulum aminopeptidase 1 (ERAP1) trimming within the ER (*18*).

As a single-stranded RNA virus, SARS-CoV-2 has a high propensity for mutation due to a lack of proof-reading ability by the RNA-dependent RNA polymerases that replicate their genomes (*19*). This has led to the emergence of several variants of concern since the start of the pandemic. Currently, the effects of SARS-CoV-2 epitope-flanking mutations on viral epitope antigen processing are unknown. Our aim was to identify whether mutations are occurring in these regions with possible effects on CD8^+^ T cell activation.

## RESULTS

### Identification of mutations in flanking regions of SARS-CoV-2 nucleoprotein CD8 epitopes

Focussing on two immunodominant SARS-CoV-2 nucleoprotein (NP) epitopes previously identified, NP_9- 17_ and NP_105-113_ (*2, 20*), we searched sequences derived from the GISAID database (www.gisaid.org; 6^th^ July 2020) for mutations within six amino acids either side of these epitopes, which yielded 29 mutations (***Fig. 1A***). Mutations found in five or fewer sequences (<10%) were excluded from further investigation to eliminate mutations that were due to sequencing artefacts, resulting in 19 mutations (***Supplementary Table 1***). We subjected these sequences through the IEDB (www.iedb.org) NetChop 20S 3.0 prediction tool for proteasomal cleavage (*21*) and compared cleavage against wild-type (WT) NP sequence derived from the viral strain originally isolated in Wuhan, China (***Fig. 1B-C***). The mutations that showed the greatest predicted differences in proteasomal cleavage as compared to WT NP sequence for each epitope were selected for further investigation (n=7) (***Fig. 1D***). While the frequency of these mutations was extremely low at the time of analysis, they have been steadily rising since July 2020 (***Fig. 1E-F***). They are not currently present in any variants of interest or concern identified by the WHO (https://www.who.int/en/activities/tracking-SARS-CoV-2-variants/).

**Figure 1:**
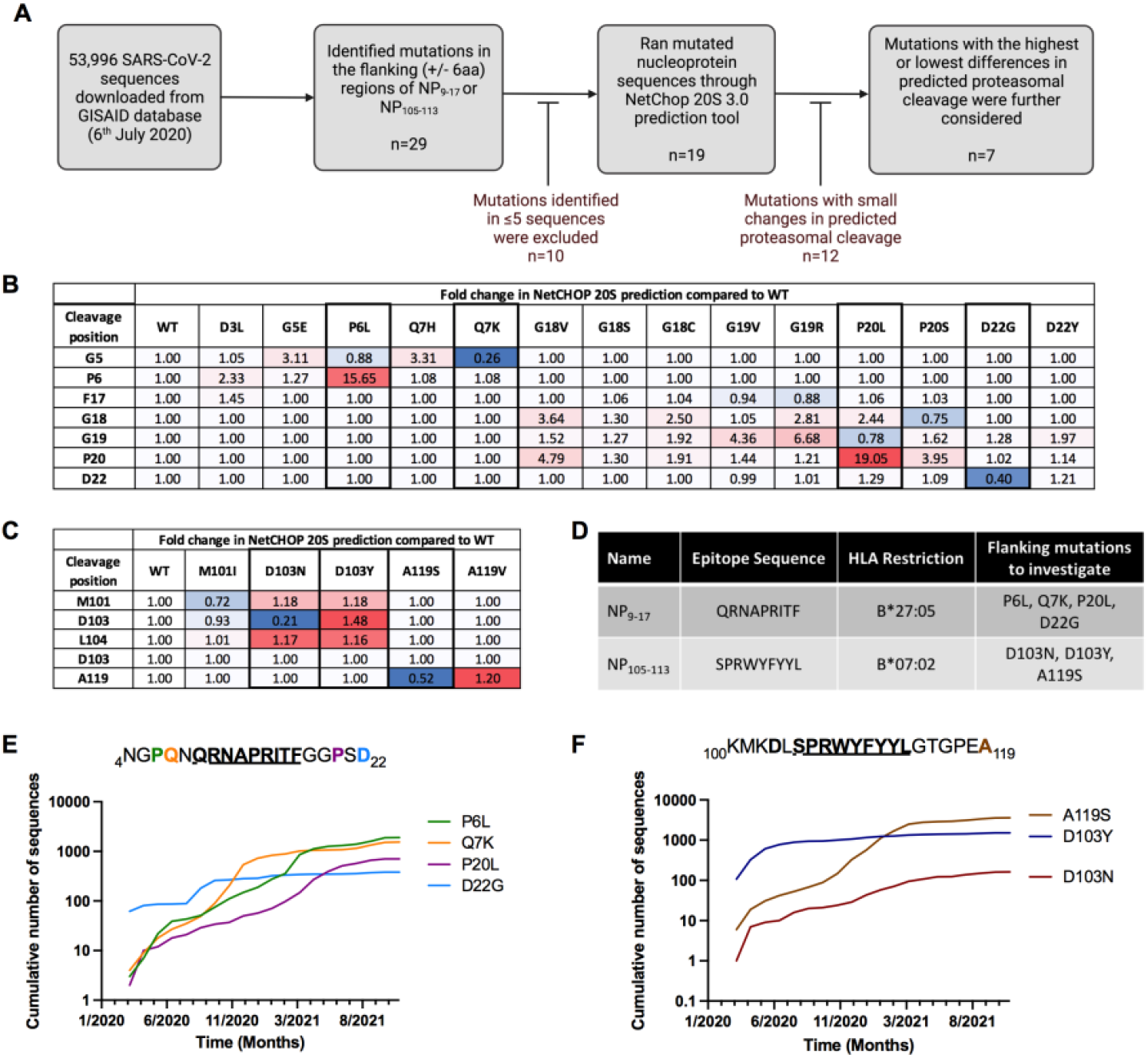
Identification of mutations in SARS-CoV-2 CD8^+^ T cell nucleoprotein epitope flanking regions. (A) 53,996 SARS-CoV-2 sequences were downloaded from the GISAID database on 6^th^ July 2020. Mutations in the flanking regions (± 6 amino acids) were assessed either side of identified CD8 nucleoprotein epitopes (NP_9-17_ / NP_105-113_) and analysed the potential effect these mutations may have on antigen processing by the proteasome. (B) Heatmap results from NetChop 20s prediction tool for the effect of mutations on proteasomal cleavage of NP_9-17_ region. Red = increase in cleavage, blue = decrease. (C) Heatmap results from NetChop 20s prediction tool for the effect of mutations on proteasomal cleavage of NP_105-113_ region. Red = increase in cleavage, blue = decrease. (D) Mutations taken forward for further investigation. (E) Frequency of mutations for NP_4-22_ region expressed as cumulative number of sequences with each mutation since January 2020. (F) Frequency of mutations for NP_100-119_ region expressed as cumulative number of sequences with each mutation since January 2020. Data downloaded from COVID-19 Viral Genome Analysis Pipeline.

### Unaltered HLA expression in SARS-CoV-2 NP-transduced B cells

Having identified mutations of interest, we next created stable B cell lines expressing SARS-CoV-2 WT or mutated NP through lentiviral transduction. B cells from patient C-COV19-005 isolated and immortalised as previously described in (*2*) were chosen for initial experiments as they expressed both HLA-B*27:05 and HLA-B*07:02, the correct MHC class I restriction for NP_9-17_ and NP_105-113_, respectively (full HLA typing for donor C-COV19-005 is available in the methods section). Each NP construct also contained a GFP gene separated by a viral self-cleavage T2A site, resulting in one long transcript but two translated proteins, GFP and NP. NP-transduced cells showed no significant differences in GFP, SARS-CoV-2 NP or HLA expression (***Supplementary Fig. 1***).

### Flanking mutations alter CD8^+^ T cell responses

Our NP-transduced B cell system allowed us to evaluate the effect of one amino acid mutation on T cell responses to NP. We recognise that this is not a complete surrogate for viral infection as NP protein expression in transduced cells may be greater than that of a SARS-CoV-2 infection. However, NP is the most abundant viral protein at all stages of the infection cycle, especially in the initial stages (*22, 23*), and a recent study showed that comparable levels of NP could be detected following virus infection as compared to transfection of HEK293T cells (*24*).

Using NP-transduced cell lines as target cells, we co-cultured with bulk CD8^+^ T cells-specific for NP_9-17_ or NP_105-113_ for four hours prior to analysis of CD107a, IFNγ and TNFα expression (***Fig. 2A***). We included the mutation NP-P13L as a control in our experiments as previous work had shown this to be a CD8^+^ T cell escape mutant within the epitope NP_9-17_ (*17*). This mutation has subsequently been found within the Omicron variant of SARS-CoV-2 (https://covariants.org/variants/21K.Omicron). As expected, the NP-P13L mutation was shown to lead to complete escape from all NP_9-17_ CD8^+^ T cell responses measured when compared to WT NP (***Fig. 2B***).

**Figure 2:**
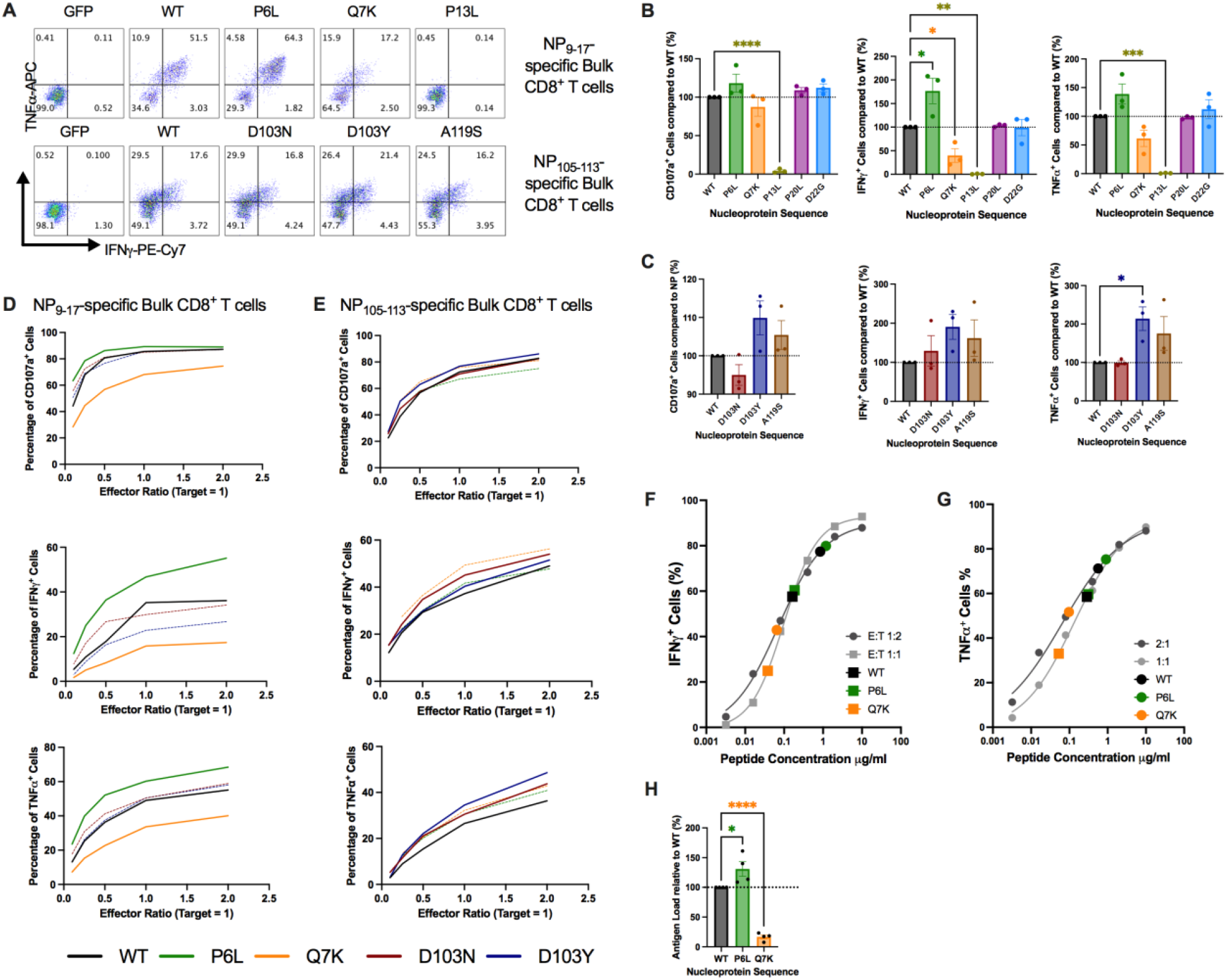
Mutations in flanking regions of epitopes alter CD8^+^ T Cell responses. (A) Intracellular cytokine staining of NP_9-17_ or NP_105-113_ epitope-specific bulk CD8^+^ T cells following incubation with GFP-or NP-transduced B cells. (B) CD8^+^ T Cell responses to mutations flanking NP_9-17_ were analysed by flow cytometry at an E:T ratio of 2:1. (C) CD8^+^ T Cell responses to mutations flanking NP_105-113_ were analysed by flow cytometry at an E:T ratio of 2:1. (D-E) Selected mutations were investigated in more detail by altering the E:T ratio from 2:1 to 0.1:1. (F) IFNγ titration curve in response to NP_9-17_ epitope peptide for NP_9-17_-specific bulk CD8^+^ T cells with NP-transduced B cell responses overlaid. (G) TNFα titration curve in response to NP_9-17_ epitope peptide for NP_9- 17_-specific bulk CD8^+^ T cells with NP-transduced B cell responses overlaid. (H) Estimated peptide load values were normalised to WT-NP levels. Data is presented as mean ±SEM and was analysed by one-way ANOVA and Dunnett’s multiple comparisons test. \**p*<0.05, \*\**p*<0.01, \*\*\**p*<0.001, \*\*\*\**p*<0.0001.

Interestingly, we also saw significant differences in T cell responses to several flanking mutations (***Fig. 2B-C***). NP-Q7K significantly decreased IFNγ^+^ NP_9-17_-specific CD8^+^ T cells (*p=0.0448*). In contrast, NP-P6L was shown to significantly increase the proportion of IFNγ^+^ NP_9-17_-specific T cells (*p=0.0107*), while NP-D103Y significantly increased TNFα^+^ NP_105-113_-specific T cells (*p=0.0424*).

Addressing the mutations that showed significant differences, in addition to NP-D103N as it affects the same amino acid position as NP-D103Y, we investigated how these mutations modulate CD8^+^ T cell responses in more detail. The previous experiment was performed at an effector (CD8^+^ T Cell) to target (NP-transduced B cell) ratio (E:T) of 2:1. We next used a range of E:T from 2:1 to 0.1:1 to expose differences in CD8^+^ T cell responses (***Fig. 2D-E***). For the mutations flanking NP_9-17_, there was clear separation between the results for WT vs mutant, with NP-P6L always showing higher and NP-Q7K showing lower T cell responses. For NP_105-113_, the differences were more subtle, but there is separation between WT and mutant proteins. Taken together, these results clearly demonstrate that flanking mutations, particularly N-terminal mutants, can significantly alter CD8^+^ T cell responses, presumably by altering the antigen load of the epitopes on the cell surface.

### Flanking mutations alter antigen load

To confirm whether the differences in CD8^+^ T cell responses were linked to cell surface antigen load, we pulsed exogenous epitope NP_9-17_ peptide over un-transduced B cells at concentrations of 0.0032-10 mg/ml for one hour prior to co-culture with NP_9-17_-specific bulk CD8^+^ T cells at both E:T 2:1 and 1:1. At the same time, we co-cultured NP_9-17_-specific bulk CD8^+^ T cells at both E:T 2:1 and 1:1 with our NP-transduced B cell lines. After four hours of co-culture, we evaluated CD8^+^ T cell activation by intracellular cytokine staining for IFNγ and TNFα. NP_9-17_ epitope was selected for further analysis as it showed the greatest differences across a range of E:T ratios (***Fig. 2D***).

Plotting the peptide titration curve for each cytokine, we were able to extrapolate an estimated peptide concentration for each NP-transduced cell line (***Fig. 2F-G***). These estimated peptide load values showed a broad range suggesting that estimation of accurate peptide concentration is not possible using this technique. However, when each peptide load value was compared to WT-NP, significant differences could be seen for both NP-P6L (*p*=0.0273) and NP-Q7K (*p*<0.0001) (***Fig. 2H***) showing that this system is valuable in comparing antigen load for mutants versus WT-NP.

### Flanking mutations alter epitope generation by the proteasome

CD8^+^ T cell epitopes are generated through degradation of proteins by the proteasome, machinery that selectively cleaves after certain amino acid residues. Therefore, we investigated whether our flanking mutations alter proteasomal digestion of NP epitopes. To do this, we digested long peptide precursors (NP_1- 22_ and NP_99-126_), containing the epitopes NP_9-17_ and NP_105-113_ and either WT or mutated flanking regions with constitutive (20S) or immunoproteasome (i20S) for up to 3 hours *in vitro*, followed by analysis of the digestion products via quantitative mass spectrometry (MS).

The immunoproteasome is constitutively expressed in haematopoietic cells or induced in response to IFNγ or TNFα and can produce antigenic peptides at a faster rate than the constitutive proteasome (*25, 26*). As a haematopoietic cell line, we expected that our B cell lines would express a combination of 20s and i20s proteasomes, which was confirmed through detection of constitutive proteasome (PSMB6) and immunoproteasome (LMP2, LMP7) subunits (***Supplementary Fig. 2***).

The products of NP_1-22_ with a 3-hour immunoproteasome digestion showed a distinct pattern of products for NP-WT, NP-P6L and NP-Q7K (***Fig. 3A***). Strikingly, for both NP_9-17_ and NP_105-113_ epitopes, proteasomal cleavage alone was sufficient to produce not only correctly C-terminally cleaved peptides, but also the complete nine amino acid (9mer) epitopes when embedded in WT flanking sequences, regardless of the type of proteasome (***Fig. 3***). This is remarkable as epitope precursors generated by the proteasome usually require additional processing, particularly at their N-terminus (*27, 28*).

**Figure 3:**
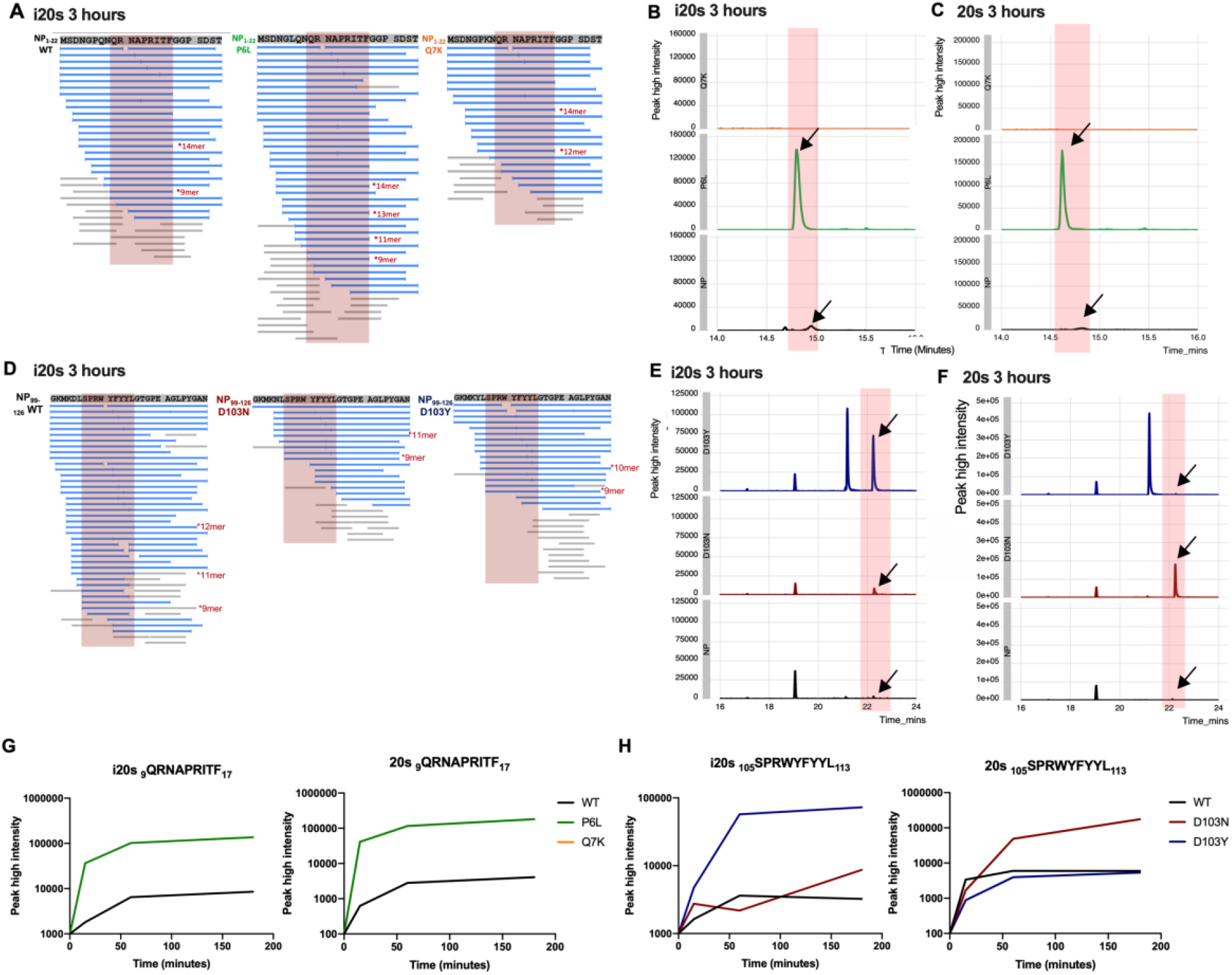
Mutations in flanking regions affect proteasomal digestion and epitope generation of SARS-CoV-2 CD8^+^ T cell epitopes. (A) PEAKS data showing the products of NP_1-22_ peptide digested with immunoproteasome for 3 hours. Correctly C-terminally cleaved peptides are labelled for 9-14mer peptides. (B) Chromatograms showing generation of NP_9-17_ epitope after 3-hour incubation of NP_1-22_ peptide with immunoproteasome. (C) Chromatograms showing generation of NP_9-17_ epitope after 3-hour incubation of NP_1-22_ peptide with constitutive proteasome. (D) PEAKS data showing the products of NP_99-126_ peptide digested with immunoproteasome for 3 hours. Correctly C-terminally cleaved peptides are labelled for 9-14mer peptides. (E) Chromatograms showing generation of NP_105-113_ epitope after 3-hour incubation of NP_99-126_ peptide with immunoproteasome. The additional peaks correspond to NP_103-110_ and NP_112-124_. (F) Chromatograms showing generation of NP_105-113_ epitope after 3-hour incubation of NP_99-126_ peptide with constitutive proteasome. The additional peaks correspond to NP_103-110_ and NP_112-124_. (G) Time-course of NP_9-17_ epitope generation from NP_1-22_ long peptide with variants digested with immuno- or constitutive proteasome. (H)Time-course of NP_105-113_ epitope generation from NP_99-126_ long peptide with variants digested with immuno- or constitutive proteasome.

Across a 3-hour time-course, we observed higher NP_9-17_ epitope generation with NP-P6L as compared to WT sequences, which was consistent with both immunoproteasome and constitutive proteasome digestions (***Fig. 3G***). We were unable to detect NP_9-17_ epitope from NP-Q7K mutated protein sequences with either proteasome (***Fig. 3A-C, 3G***). We identified proteasomal differences in the response to mutated proteins in the NP_99-126_ peptide, with NP-D103N increasing NP_105-113_ epitope generation by the constitutive proteasome but having an insignificant effect with the immunoproteasome. In contrast, NP-D103Y enhanced NP_105-113_ epitope generation by immunoproteasome digestion (***Fig. 3H***).

### Flanking mutations alter N-terminally extended proteasomal products

In addition to differences in 9mer epitope generation by the proteasome, we found that mutations also affected generation of N-terminally extended epitope precursors (***Fig. 4***). This appears to be particularly prevalent for the mutations NP-P6L (***Fig. 4A-B***) and NP-D103N (***Fig. 4C*)**. We also observed differences dependent on proteasome species (***Fig. 4C-D***). N-terminally extended precursors are most commonly trimmed by endoplasmic reticulum aminopeptidase 1 (ERAP1) in the ER to the optimal length for HLA Class I binding (*29*), confirmed to be nine amino acids for NP_9-17_ and NP_105-113_ (*2*). The requirement for further processing of epitopes can alter the efficiency of antigen processing and presentation potentially affecting antigen load at the cell surface.

**Figure 4:**
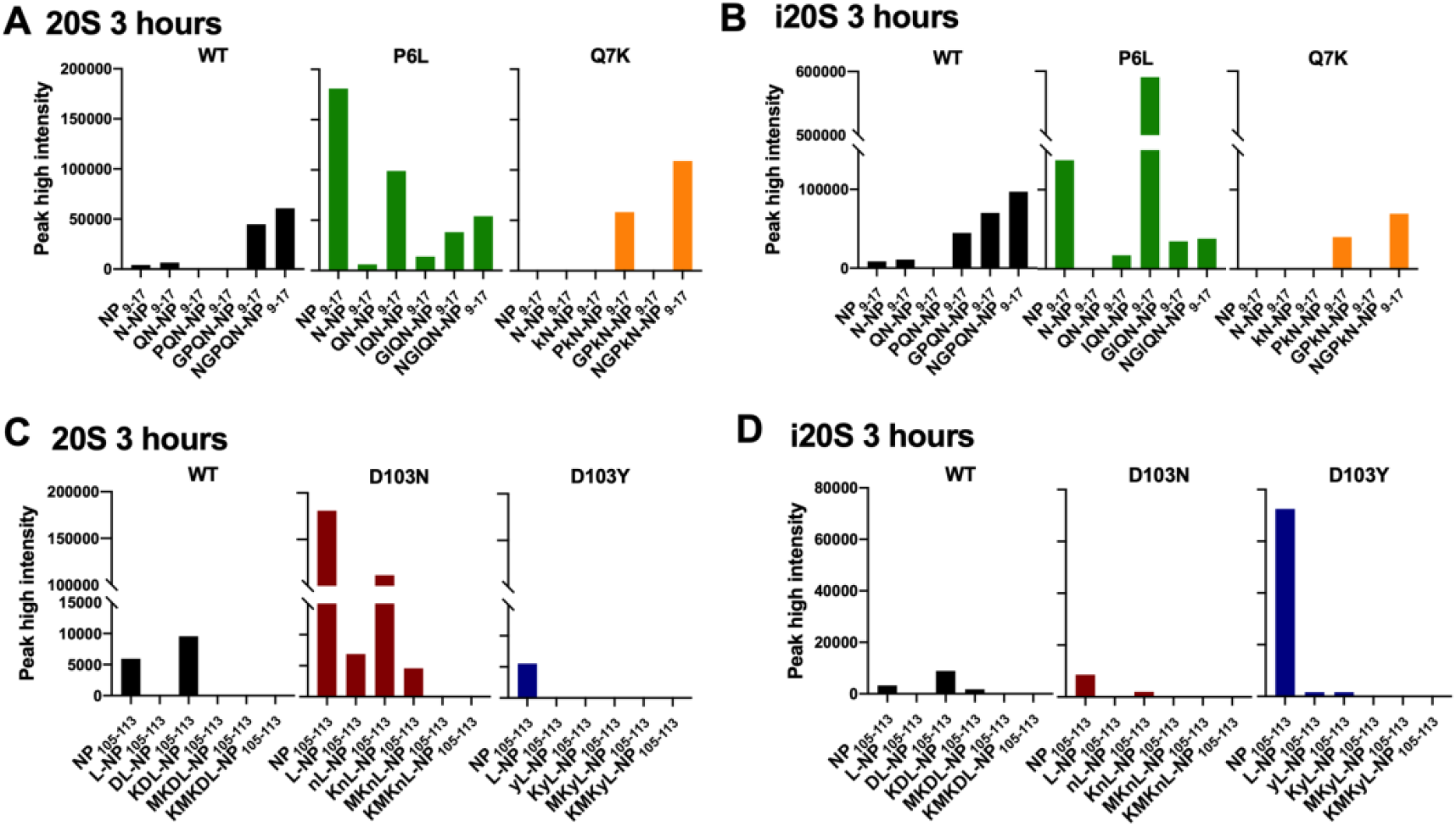
Flanking mutations alter N-terminally extended proteasomal products. (A) The generation of 9-14mers (9mer = NP_9-17_) from NP_1-22_ peptide following digestion with constitutive proteasome for 3 hours. (B) The generation of 9-14mers (9mer = NP_9-17_) from NP_1-22_ peptide following digestion with immunoproteasome for 3 hours. (C) The generation of 9-14mers (9mer = NP_105-113_) from NP_99-126_ peptide following digestion with constitutive proteasome for 3 hours. (D) The generation of 9-14mers (9mer = NP_105-113_) from NP_99-126_ peptide following digestion with immunoproteasome for 3 hours.

## DISCUSSION

For SARS-CoV-2 infection, it has been demonstrated that mutations within epitope sequences can lead to escape from T cell recognition (*16*), presumably by disruption of HLA-TCR engagement. The effect of mutations within the flanking regions of SARS-CoV-2 epitopes has yet to be established. Here, we show that mutations in these regions do in fact influence CD8^+^ T cell responses due to differences in the processing of viral antigens. Most importantly, we show that mutations such as NP-Q7K are capable of decreasing T cell responses and potentially leading to escape through reductions in cell surface antigen load because of poor proteasomal processing of the epitope.

We established a system which enabled us to evaluate T cell responses to naturally processed epitope peptides and to natural mutations that impact the delivery of epitope antigen to the cell surface even when the epitope sequence is conserved. We, and others (*30*), believe that continual monitoring of potential escape from immunodominant T cell responses in circulating variants before they become dominant strains is critically important for COVID-19 surveillance and future vaccine strategies.

We showed that both NP-P6L and NP-D103Y could increase CD8^+^ T cell responses due to increased efficiency of proteasomal digestion and potentially higher epitope peptide load on the cell surface, suggesting that some variants identified in the NP sequence could be beneficial for viral fitness, representing an advantage over increasing CD8^+^ T cell responses. We believe continued identification of such “beneficial” natural mutations within immunodominant T cell epitope flanking regions could provide a new avenue of research for vaccine design, especially as the inclusion of NP into future vaccine designs against SARS-CoV-2 shows promise (*31, 32*).

The number of mutations investigated in this study reflects a small subset due to the early timepoint of sequence sampling, but three out of the twenty-nine mutations (∼10%) originally identified showed significant differences in CD8^+^ T cell responses. Moreover, four mutations (∼14%) investigated for their effect on proteasomal digestion showed differences in the amount of 9mer epitope peptide generated. This suggests that there are many mutations that may be naturally occurring in the flanking regions of epitopes that have the potential to alter T cell responses.

Our results emphasise that the antiviral efficacy and cross-reactive potential of CD8^+^ T cells should be assessed in response to virus-infected cells, or minimally within systems that allow natural processing of epitopes. Tracking and characterisation of epitope-flanking regions in a system where natural processing of viral antigens occurs will allow us to make more precise judgements of the risk of future viral strains.

## MATERIALS AND METHODS

### Identification of mutations of interest proximal to SARS-CoV-2 CD8 nucleoprotein epitopes

On 6^th^ July 2020, all available sequences for SARS-CoV-2 that differed from the original Wuhan strain were downloaded from the GISAID database (www.gisaid.org). The regions NP_4-22_ and NP_100-119_ were analysed for mutations. Those falling within the epitope sequence region or found in less than 5 sequences were discounted for further analysis (***Supplementary Table 1***).

### Prediction tool analysis of proteasomal digestion

Using the NetChop 20S 3.0 prediction tool (www.iedb.org) (*21*), we evaluated the predicted proteasomal cleavage sites for each peptide region in both WT and mutated sequences (***NP***_***4-22***_**: *Supplementary Table 2, NP***_***100-119***_**: *Supplementary table 3***). Data was presented as fold-change compared to the predicted proteasomal cleavage in WT sequences (***Fig. 1B-C***).

### Tracking of mutation frequencies

The frequency of mutations over time was assessed from data downloaded from the COVID-19 Viral Genome Analysis Pipeline (https://cov.lanl.gov/content/index) tracking mutations tool (*33*). Data was accessed on 10^th^ December 2021.

### Lentivirus construction & production

SARS-CoV-2 nucleoprotein (NP) was cloned from pHRSIN (kind gift from Prof. Alain Townsend) into the Addgene 17448 lentivirus vector by InFusion PCR (TaKaRa) and verified by Sanger sequencing. NP expression was coupled to eGFP expression using a T2A linker. Single nucleotide mutations were introduced using site-directed mutagenesis (SDM) (New England BioLabs, NEB Base Changer kit). Once confirmed, mutated gene cassettes were extracted by restriction digest and cloned into a clean vector to prevent any off-target effects of SDM-PCR.

Lentivirus was produced using HEK293T cells, expression constructs were co-transfected with psPAX2 and pMD2.G using PEIPro. Lentiviral supernatants were filtered using a 0.45µm CA filter and overlayed on 20% sucrose before ultracentrifugation at 90,000g for 120 minutes at 4°C. Pelleted virus was resuspended in PBS, aliquoted and stored at -80°C.

### Lentivirus transduction

Cells from donor C-COV19-005 were used for investigation as they contained the HLA for both NP_9-17_-HLA-B*27:05 and NP_105-113_-HLA-B*07:02. Full HLA Class I type for this donor is: A*01:01, A*02:01, B*07:02, B*27:05, C*01:02 and C*07:02. This EBV-transformed B cell line generated from patient samples as previously described in (*2*), was spin inoculated with lentivirus to generate stable cell lines. In brief, B cells were plated at 5×10^5^ cells per well of a 96-well U-bottom plate and supplemented with 5×10^6^ TU/ml lentivirus and 8mg/ml polybrene in RPMI + 1% FBS. The plate was spun at 500g, 37°C for 1 hour prior to overnight incubation at 37°C. After 24 hours, B cells were washed and cultured in RPMI + 10% FBS until enough cells were obtained for cell sorting. Cells were then bulk sorted for GFP expression on a BD FACSAria III Cell sorter.

### Cell culture of transduced B cell lines & T cells

B cell lines were cultured in RPMI (Lonza) + 10% foetal calf serum (Sigma) + 2mM L-glutamine (Sigma) at 37°C, 5% CO_2_. Bulk T cells generated as previously described in (*2, 20*) were cultured in RPMI (Gibco) + 10% human serum (Sigma) + 2mM L-glutamine (Sigma) + Penicillin-Streptomycin (Sigma, 100 units penicillin and 100 ug streptomycin per mL), supplemented with 100 U/ml IL-2 (Sigma).

### Flow cytometry analysis of transduced B cells

Transduced and sorted B cell lines were stained with Zombie-Violet live dead stain (Thermofisher) and PE-anti-HLA-ABC (Biolegend) and run on an Attune NXT Flow cytometer (Thermofisher). Data was analysed using FlowJo v10 (BD Biosciences).

### Western blot analysis of transduced B cells

Transduced B cell lines were homogenized using RIPA lysis buffer (Thermo Fisher Scientific). Anti-glyceraldehyde 3-phosphate dehydrogenase (GAPDH) antibody clone 6C5 (Merck Millipore, MAB374) was used as control antibody; anti-SARS-CoV-2 nucleocapsid antibody (2µg/µl, Sino Biological, 40143-R001) was used to probe NP expression. Anti-PSBM6 (AbCam), anti-LMP2 (AbCam) and anti-LMP7 (AbCam) were used individually at 1 in 1000 dilution to assess proteasomal subunit expression in the transduced B cells.

Primary antibodies were probed by IRDye 680LT goat anti-mouse (Li-Cor, 926-68020) and IRDye 800LT goat anti-rabbit (Li-Cor, 925-32210), and visualized using the iBright FL1000 (Invitrogen). Images from Western blot experiments were analysed by Fiji software for band density and expressed in GraphPad Prism as a percentage of the GAPDH expression. A one-way ANOVA was performed along with Tukey’s or Sidak’s multiple comparisons tests to measure statistical differences between cell lines.

### Intracellular cytokine staining of T cells

Bulk CD8^+^ T cells specific for NP_9-17_ or NP_105-113_ epitope were co-cultured with transduced B cell lines for 4 h with BD GolgiPlug, BD GolgiStop and PE-anti-CD107a. Dead cells were labelled using Fixable Zombie Violet dye from Thermofisher. Cells were then washed, fixed with Cytofix/Cytoperm (BD Biosciences), and stained with PE-Cy7-anti-IFNγ (eBioscience), APC-anti-TNFα (eBioscience) and BV711-anti-CD8 (Biolegend). Cells were fixed with BD CellFix and run on an Attune NXT Flow Cytometer (Thermofisher).

For analysis, live CD8^+^ T cells were gated on and analysed for their expression of CD107a, IFNγ or TNFα using FlowJo V10 (BD Biosciences). Using each bulk T cell line as a control for the other we calculated the percentage response of NP_9-17_ over NP_105-113_, and vice versa. This data was then normalised to a percentage of the response to WT NP. Data was plotted and analysed in GraphPad Prism v9.2.0. Statistical analysis of responses was done by one-way ANOVA and Dunnett’s multiple comparisons test.

### Determining epitope antigen load

WT donor C-COV19-005 B cells were pulsed with exogenous NP_9-17_ epitope peptide ranging in concentration from 0.0032-10 mg/ml for 1 hour prior to co-culture with bulk NP_9-17_ CD8^+^ T cells for 4 hours and intracellular cytokine staining (ICS) as described above. Transduced B cell lines were also co-cultured with bulk NP_9-17_ CD8^+^ T cells for 4 hours prior to ICS.

Data was plotted and analysed in GraphPad Prism v9.2.0. The percentage of IFNγ or TNFα cells were plotted on individual x/y graphs and fit with a non-linear curve. For transduced cells, the data was also plotted along the curve based on the percentage of cytokine^+^ cells. Using the non-linear curve data, estimated peptide concentrations were extrapolated for each transduced B cell line. These peptide load values were then normalised to a percentage of NP-transduced cells and presented in a column graph. Data was analysed by one-way ANOVA comparison and Dunnett’s multiple comparison’s test comparing mutant transduced cells to WT NP-transduced cells.

### Statistical Analysis of CD8^+^ T cell responses

Experiments were repeated to an *n* of 3. All data with statistical analysis was analysed by one-way ANOVA using Dunnett’s multiple comparisons test. ANOVA summaries: NP_9-17_ CD107a – F = 33.95, P value <0.0001; NP_9-17_ IFNγ -F = 17.72, P value <0.0001; NP_9-17_ TNFα - F = 17.87, P value <0.0001; NP_105-113_ CD107a – F = 4.095, P value = 0.0492; NP_105-113_ IFNγ - F = 1.323, P value = 0.3328; NP_105-113_ TNFα - F = 4.492, P value = 0.0397. Brown-Forsythe tests: NP_9-17_ CD107a – F (DFn, DFd) = 0.6481 (5,12), P value = 0.6685; NP_9-17_ IFNγ - F (DFn, DFd) = 0.8037 (5,12), P value = 0.5683; NP_9-17_ TNFα - F (DFn, DFd) = 1.382 (5,12), P value = 0.2981; NP_105-113_ CD107a – F (DFn, DFd) = 0.5568 (3,8), P value 0.6581; NP_105-113_ IFNγ - F (DFn, DFd) = 0.5392 (3,8), P value = 0.6686; NP_105-113_ TNFα - F (DFn, DFd) = 0.9408 (3,8), P value = 0.4650.

### Proteasomal digestion of peptides

Peptides were purchased from GenScript (information available in **Supplementary Table 4**) and reconstituted to 20mg/ml in DMSO. Recombinant human proteasomes (Human 20S Proteasome Protein, E360-050, bio-techne; Human 20S Immunoproteasome Protein, E370-025, bio-techne) and peptides were combined at a ratio of 50:1 for a period of 15, 60 or 180 minutes prior to the addition of 0.1µl formic acid to terminate the function of the proteasome. Incubations were carried out at 37°C with rotation to increase proteasome activity. Digested samples were frozen and stored at -80°C until analysis by mass spectrometry.

### Analysis of in vitro proteasome antigen processing products by mass spectrometry

Products generated by *in vitro* proteasome digestion of antigen precursor peptides were analysed by nano-flow liquid chromatography tandem mass spectrometry (nLC-MS/MS) using a Dionex Ultimate 3000 (Thermo) coupled to a timsTOF Pro trapped ion mobility spectrometry (TIMS) time-of-flight mass spectrometer (Bruker) using a 75 µm × 150 mm C18 column with 1.6 µm particles (IonOpticks) at a flow rate of 400 nL/min at 40 °C. The gradient used was a 17-minute increase from 2 % buffer B to 30 % buffer B, followed by an increase to 95 % B in 30 seconds and a 2.5-minute wash and 5 minutes re-equilibration at 2 % B (A: 0.1 % formic acid in water. B: 0.1 % formic acid in acetonitrile).

The timsTOF Pro mass spectrometer was operated in data dependent acquisition - parallel accumulation, serial fragmentation (ddaPASEF) mode. The TIMS accumulation and ramp times were set to 100 ms, and mass spectra were recorded from 100 – 1700 m/z, with a 0.85 – 1.30 Vs/cm^2^ ion mobility range. Precursors were selected for fragmentation from an area of the full TIMS-MS scan that excludes the majority of ions with a charge state of 1+. Those selected precursors were isolated with an ion mobility dependent collision energy, increasing linearly from 27 – 45 eV over the ion mobility range. Four PASEF MS/MS scans were collected per full TIMS-MS scan using a 3-cycle overlap, giving a duty cycle of 0.53 s. Ions were included in the PASEF MS/MS scan if they met the target intensity threshold of 2000 and were sampled multiple times until a summed target intensity of 10000 was reached. A dynamic exclusion window of 0.015 m/z by 0.015 Vs/cm^2^ was used, and sampled ions were excluded from reanalysis for 24 seconds.

Raw MS data files were searched against a database containing the full synthetic antigen peptide precursor sequences using Peaks Studio Version 10 with mass tolerances of 20.0 ppm for precursors and 0.04 Da for fragment ions. Nonspecific cleavage was used and oxidation and deamidation were allowed as variable modifications. Peptide abundance was measured as accumulated ion counts after extraction of the chromatographic peak profiles of the precursor masses using Skyline version 21 and Bruker Compass DataAnalysis Version 5.3. Chromatograms were generated using R software package v4.0.1 and ggplot2 v3.3.2 and ggridges v0.5.2. The mass spectrometry proteomics data have been deposited to the ProteomeXchange Consortium via the PRIDE (*34*) partner repository with the dataset identifier PXD032054.

## Supplementary Materials

**Supplementary Figure 1:**
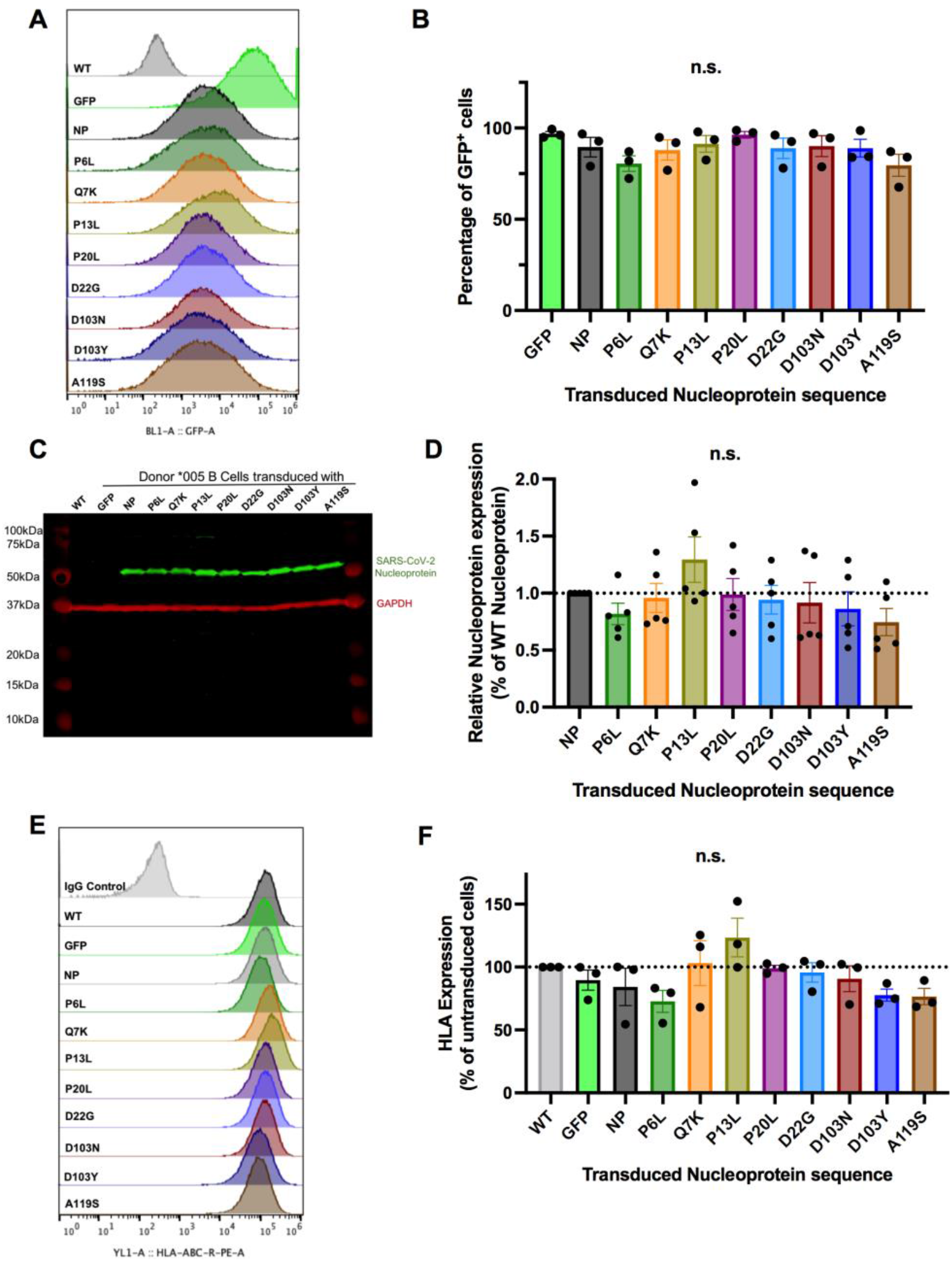
Characterisation of nucleoprotein-transduced B Cells. (A) Histograms showing GFP expression of transduced C-COV19-005 B cells following cell sorting. (B) The percentage of GFP^+^ cells per cell line. (C) Representative western blot for SARS-CoV-2 Nucleoprotein (NP) and GAPDH expression. D) Western blot results normalised to GAPDH expression and shown as a percentage of the NP expression of wild-type (WT) NP. (E) Histograms showing HLA-ABC expression on WT and transduced #005 B cells. (F) The HLA expression on transduced cells shown as a percentage of the HLA expression on WT un-transduced cells. For all experiments, n=3. Data analysed by one-way ANOVA and Dunnett’s multiple comparisons test.

**Supplementary Figure 2:**
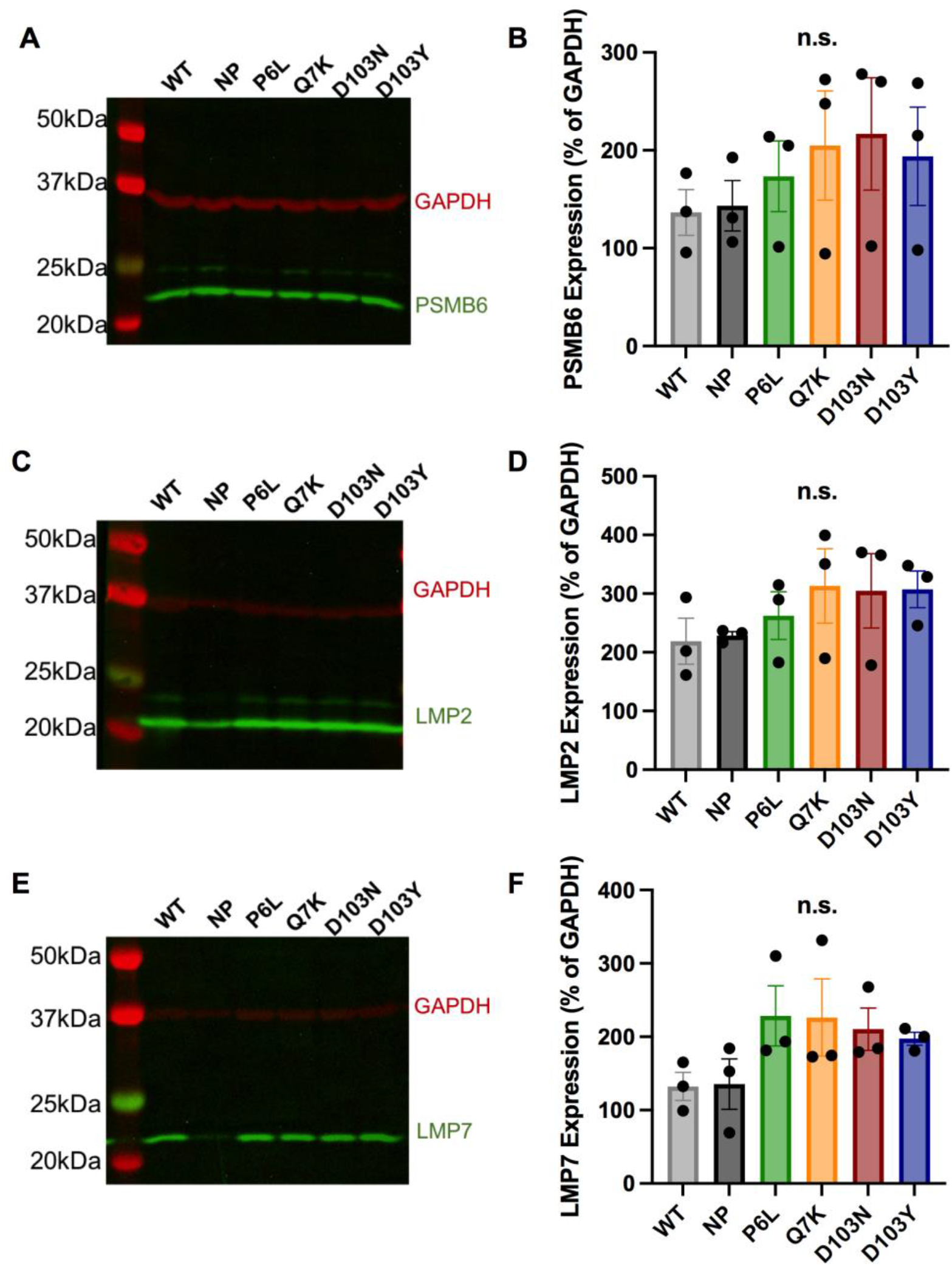
Proteasomal subunit expression in B cell lines. (A) The constitutive proteasomal subunit PSMB6 was detected in B cell lines by western blot in concert with GAPDH. (B) Expression of PSMB6 as a percentage of GAPDH is shown for each cell line. (C) The immunoproteasomal subunit LMP2 was detected in B cell lines by western blot in concert with GAPDH. (D) Expression of LMP2 as a percentage of GAPDH is shown for each cell line. (E) The immunoproteasomal subunit LMP7 was detected in B cell lines by western blot in concert with GAPDH. (F) Expression of LMP7 as a percentage of GAPDH is shown for each cell line. For each experiment, n=3. Data analysed by one-way ANOVA and Dunnett’s multiple comparisons test.

**Supplementary Table 1:** Identified mutations in flanking regions of NP_9-17_ and NP_105-113_ and their frequencies as of July 2020 and December 2021.

| Epitope Name | Flanking mutation | July 2020 Frequency |  | December 2021 Frequency |  | Change in percentage |
| --- | --- | --- | --- | --- | --- | --- |
|  |  | Number | Percentage | Number | Percentage |  |
| NP9-17 | D3L | 13 | 0.0241 | 697,514 | 31.4037 | + 31.3796% |
|  | G5E | 11 | 0.0204 | 489 | 0.0220 | + 0.0016% |
|  | P6L | 36 | 0.0667 | 2,319 | 0.1043 | + 0.0376% |
|  | Q7H | 21 | 0.0389 | 175 | 0.0079 | - 0.031% |
|  | Q7K | 13 | 0.0241 | 2,594 | 0.1166 | + 0.0925% |
|  | G18V | 16 | 0.0296 | 12,553 | 0.5644 | + 0.5348% |
|  | G18S | 8 | 0.0148 | 286 | 0.0129 | - 0.0019% |
|  | G18C | 38 | 0.0704 | 1,211 | 0.0544 | - 0.016% |
|  | G19V | 8 | 0.0148 | 71 | 0.0032 | - 0.0116% |
|  | G19R | 7 | 0.0130 | 336 | 0.0151 | + 0.0021% |
|  | P20L | 11 | 0.0204 | 759 | 0.0341 | + 0.0137% |
|  | P20S | 8 | 0.0148 | 182 | 0.0082 | - 0.0066% |
|  | D22G | 146 | 0.2704 | 401 | 0.0180 | - 0.2524% |
|  | D22Y | 6 | 0.0111 | 623 | 0.0280 | + 0.0169% |
| NP105-113 | M101I | 6 | 0.0111 | 413 | 0.0186 | + 0.0075% |
|  | D103N | 11 | 0.0204 | 195 | 0.0088 | - 0.0116% |
|  | D103Y | 1,257 | 2.3280 | 1,630 | 0.0733 | - 2.2547% |
|  | A119S | 60 | 0.1111 | 5,798 | 0.2607 | + 0.1497% |
|  | A119V | 28 | 0.0519 | 93 | 0.0042 | + 0.08092% |

**Supplementary Table 2:** NetChop 20S prediction values for each mutation flanking NP_9-17_.

|  | NetCHOP - 20S Output |  |  |  |  |  |  |  |  |  |  |  |  |  |  |
| --- | --- | --- | --- | --- | --- | --- | --- | --- | --- | --- | --- | --- | --- | --- | --- |
| Cleavage position | WT | D3 -> L | G5 -> E | P6 -> L | Q7 -> H | Q7 -> K | G18 -> V | G18 -> S | G18 -> C | G19 -> V | G19 -> R | P20 -> L | P20 -> S | D22 -> G | D22 -> Y |
| **G5** | 0.183 | 0.836 | 0.57 | 0.161 | 0.606 | 0.047 | 0.183 | 0.183 | 0.183 | 0.183 | 0.183 | 0.183 | 0.183 | 0.183 | 0.183 |
| **P6** | 0.037 | 0.426 | 0.047 | 0.579 | 0.04 | 0.04 | 0.037 | 0.037 | 0.037 | 0.037 | 0.037 | 0.037 | 0.037 | 0.037 | 0.037 |
| **F17** | 0.909 | 0.054 | 0.909 | 0.909 | 0.909 | 0.909 | 0.912 | 0.966 | 0.944 | 0.855 | 0.802 | 0.967 | 0.932 | 0.909 | 0.909 |
| **G18** | 0.121 | 0.909 | 0.121 | 0.121 | 0.121 | 0.121 | 0.44 | 0.157 | 0.303 | 0.127 | 0.34 | 0.295 | 0.091 | 0.121 | 0.121 |
| **G19** | 0.112 | 0.121 | 0.112 | 0.112 | 0.112 | 0.112 | 0.17 | 0.142 | 0.215 | 0.488 | 0.748 | 0.087 | 0.181 | 0.143 | 0.221 |
| **P20** | 0.043 | 0.043 | 0.043 | 0.043 | 0.043 | 0.043 | 0.206 | 0.056 | 0.082 | 0.062 | 0.052 | 0.819 | 0.17 | 0.044 | 0.049 |
| **D22** | 0.728 | 0.728 | 0.728 | 0.728 | 0.728 | 0.728 | 0.728 | 0.728 | 0.728 | 0.724 | 0.735 | 0.941 | 0.791 | 0.294 | 0.88 |

**Supplementary Table 3:** NetChop 20S prediction values for each mutation flanking NP_105-113_.

|  | NetCHOP - 20S Output |  |  |  |  |  |
| --- | --- | --- | --- | --- | --- | --- |
| Cleavage position | WT | M101 -> I | D103 -> Y | D103 -> N | A119 -> S | A119 -> V |
| **M101** | 0.6546 | 0.4737 | 0.7757 | 0.7694 | 0.6546 | 0.6546 |
| **D103** | 0.344 | 0.32 | 0.5077 | 0.0733 | 0.344 | 0.344 |
| **L104** | 0.8087 | 0.8163 | 0.9398 | 0.9482 | 0.8087 | 0.8087 |
| **L113** | 0.9457 | 0.9457 | 0.9457 | 0.9457 | 0.9457 | 0.9457 |
| **A119** | 0.2849 | 0.2849 | 0.2849 | 0.2849 | 0.1489 | 0.3427 |

**Supplementary Table 4:**
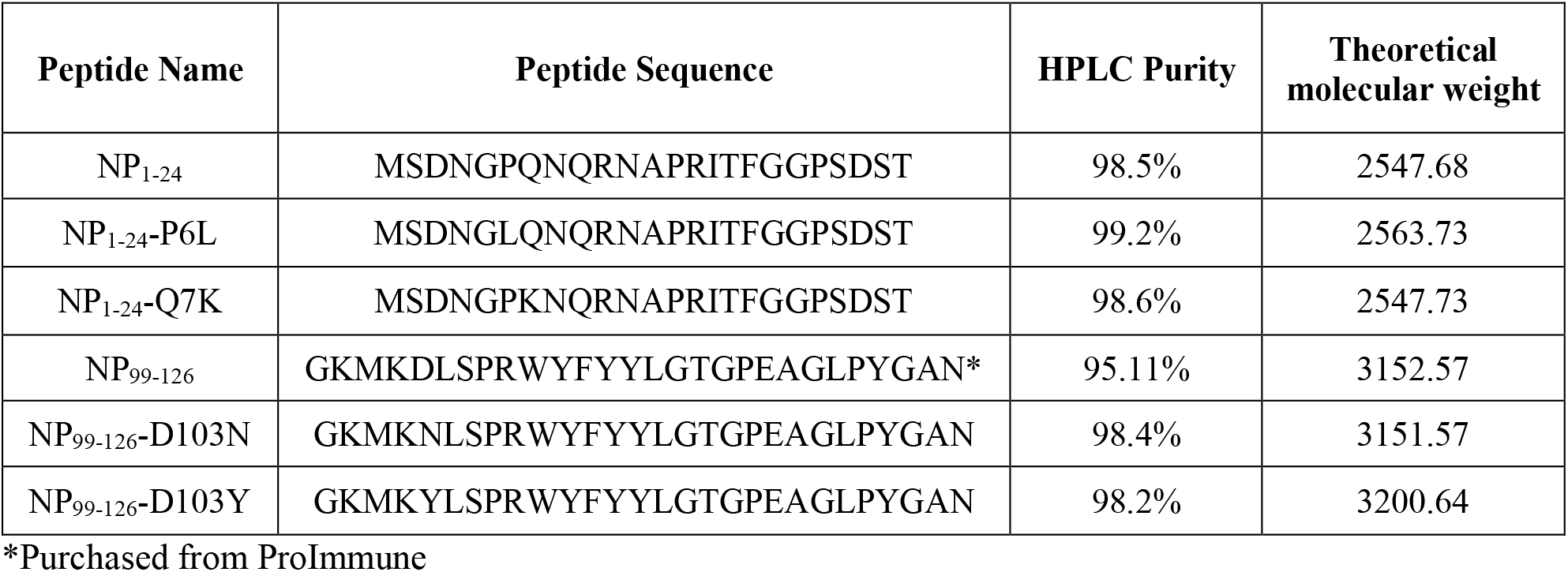
Long peptides purchased from GenScript or ProImmune for proteasomal digestion.

## Acknowledgments

We thank members of the Dong and Kessler groups for helpful discussions. We thank Prof. Alain Townsend and his group for providing valuable reagents for this work.

## Author contributions

Conceptualization: DW, BMK, TD

Wet lab work: DW, ZY, PH, RB, DD, GL, YP, XY

Mass Spectrometry: ZYu, RH, SD, RF, BMK

Data Analysis: DW, SLF

Supervision: DW, BMK, TD

Writing – original draft: DW

Writing – review & editing: DW, BMK, TD

## Data and materials availability

The mass spectrometry proteomics data have been deposited to the ProteomeXchange Consortium via the PRIDE (*34*) partner repository with the dataset identifier PXD032054. All other data are available in the main text or the supplementary materials.

